# Distinct Patterns of Cortical Reorganisation Accompany Behaviour-Specific Development in Freely Behaving Infants

**DOI:** 10.64898/2026.10.03.756058

**Authors:** Candace J. Burke, Tyler J. de Zeeuw, Denver Grimm, Nisa Hofer, Amira Jensen, Connor Mullin, Jenni M. Karl

**Affiliations:** Department of Psychology, Thompson Rivers University; Department of Data Science, Thompson Rivers University

**Keywords:** functional near-infrared spectroscopy (fNIRS), infant development, naturalistic behaviour, sensorimotor development, cortical development

## Abstract

A central premise of developmental neuroscience is that mature behaviour emerges through reciprocal interactions between infants’ spontaneous actions, neural activity, and environmental experience. Yet, much human neuroimaging research relies on highly controlled, repetitive paradigms designed to minimize behavioural variability, obscuring the very brain-behaviour dynamics that drive developmental change. Here, we introduce a naturalistic functional near-infrared spectroscopy (fNIRS) paradigm that exploits, rather than suppresses, natural variability in infants’ behaviour to investigate cortical changes associated with behavioural development. Continuous video coding of spontaneous infant behaviour was combined with finite impulse response (FIR) modelling to estimate behaviour-specific hemodynamic responses associated with reach-to-grasp, hand-to-mouth, and vocalization behaviours while accounting for temporally overlapping hemodynamic responses by treating extraneous behaviours as nuisance regressors. From 6 to 13 months of age, reach-to-grasp and vocalizations increased in frequency, while all three behaviours shortened in duration. Critically, developmental changes in cortical activation were behaviour-specific. During reach-to-grasp movements, bilateral posterior parietal activity decreased with age. In contrast, during hand-to-mouth and vocalizations, somatomotor and premotor activity increased with age. Cortical activity was also related to individual variation in the frequency and duration of these behaviours, indicating that cortical recruitment remained coupled to behavioural variation independent of age. These findings reveal behaviour-specific patterns of cortical reorganisation during infant development and establish a methodological framework for linking spontaneous behaviour to cortical activity in freely behaving infants. More broadly, this approach enables direct investigation of cortical development as it unfolds through naturalistic interactions among behaviour, neural activity, and experience.

**Significance Statement:** Continuous self-generated interactions with the environment play a critical role in driving infant brain development, yet neuroimaging research often examines infants while sleeping or in highly constrained and repetitive tasks. Here, we show that spontaneous infant behaviour can instead be leveraged to provide the temporal variability needed to resolve behaviour-specific cortical responses in freely behaving infants. By combining continuous behavioural coding with data-driven finite impulse response (FIR) modelling of overlapping hemodynamic responses, we identified behaviour-specific developmental differences in cortical recruitment without tightly constraining the infant’s behaviour. This novel approach expands developmental neuroimaging beyond sleeping or trial-based paradigms and demonstrates how naturally occurring behavioural variability can become a useful feature of fNIRS experimental design. More broadly, it generates new opportunities to directly test theories of neurobehavioural development in more ecologically valid settings.

## Introduction

A newborn baby’s seemingly random movements gradually reorganise into mature intentional behaviours (1), a transformation that must be supported by corresponding changes in developing neural networks (2, 3). Developmental theories, such as Dynamic Systems Theory (DST) and the Theory of Neuronal Group Selection (TNGS; (4)) posit that spontaneous interactions between the infant’s own body, nervous system, and the environment critically drive this neural and behavioural reorganisation (5). Yet, such theories remain largely untested in humans as doing so requires measuring neural activity in freely behaving infants – a most difficult task (6).

Functional near-infrared spectroscopy (fNIRS) seems to offer a solution as it uses non-invasive portable technology to quantify changes in oxygenated (HbO) and deoxygenated (HbR) hemoglobin in superficial cortical layers as a proxy for neural activity (7). However, it is still frequently paired with conventional neuroimaging paradigms designed for participants restrained in large brain scanners. These paradigms are built on a logic of control: constrain movement, repeatedly elicit the same action with alternating baseline rest periods, and average the hemodynamic response (HR) across trials for statistical comparison with a canonical HR function featuring a stereotypical increase in HbO and decrease in HbR lasting approximately 15-20s. In essence, natural behavioural variability is intentionally eliminated to prevent closely timed movements from generating overlapping HRs that are difficult to isolate. It is, however, nearly impossible to constrain infant behaviour in this way, and even when attempted (6), infant HRs tend to diverge from the canonical HR shape (8-10). Moreover, imposing such control eliminates the very natural movement variability that developmental theories identify as the key driver of neural and behavioural change. Thus, conventional neuroimaging paradigms based on control logic are poorly suited for testing theories of developmental neuroscience.

To address this, we inverted the conventional control logic by instead exploiting infants’ natural movement variability and using it as statistical leverage within a data-driven analytical approach. We collected synchronous behavioural and fNIRS data from 48 infants aged 6 – 13 months (116 total sessions) as they performed spontaneous reach-to-grasp movements and played freely with experimenters and caregivers. The infants’ natural movement variability functioned like an endogenous “jitter” analogous to that experimentally imposed in some functional magnetic resonance imaging (fMRI) studies (11). Because different movements occurred at irregular intervals, their associated HRs overlapped differently across time with relatively low collinearity, providing the temporal variation needed to disentangle their individual contributions to the total measured hemodynamic signal. We then used a finite impulse response (FIR) model (12, 13) to estimate the actual time course of individual HRs associated with specific movements manually coded from video. These movements served as either target or nuisance event regressors in the FIR model, enabling estimation of HRs associated with individual reach-to-grasp, hand-to-mouth, and vocalization behaviours while accounting for overlapping responses from temporally adjacent movements. Rather than assuming a prespecified HR profile for each behaviour, the FIR model estimated the time course of behaviour-specific HRs directly from the data, as demonstrated in *Figure 1*. We then interpreted the observed brain-behaviour relationships in the context of established developmental findings and proposed neural mechanisms from the existing literature. This novel approach links spontaneous infant movements directly to corresponding cortical activity enabling more ecologically valid investigation of the neural mechanisms underlying early sensorimotor development in infants.

**Figure 1.**
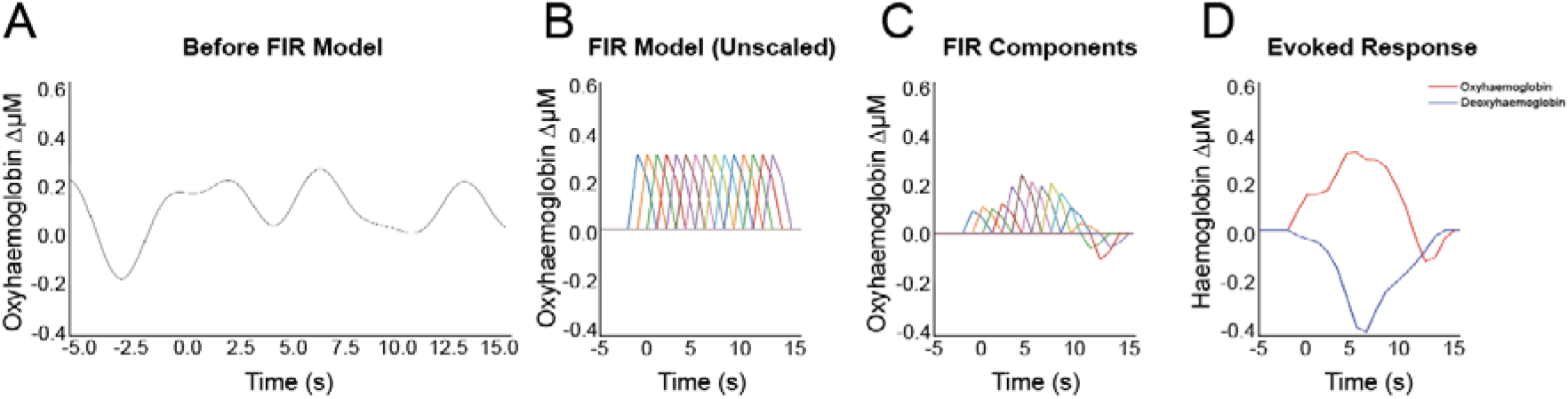
Example of finite impulse response modelling of the haemodynamic response for right-handed reach-to-grasp behaviour. Data are shown from one infant and one channel located over the right posterior parietal cortex. (A) The average oxyhaemoglobin response during right-handed reach-to-grasp events prior to modelling. (B) The unscaled FIR model, illustrating how the response window is divided across successive timepoints to estimate the haemodynamic response without assuming a predefined shape. (C) The estimated FIR components, reflecting the contribution of each successive timepoint after fitting the model to the observed data while controlling for timing and overlap of other behaviours. (D) The resulting estimated haemodynamic response during right-handed reach-to-grasp behaviour, shown for oxyhaemoglobin (red) and deoxyhaemoglobin (blue).

## Results

### 2.1. Behaviour-Specific Changes Across Age

With increasing age, the frequency of right-handed reaches *(Supplemental Table 1)* increased (β = 0.087, *p* = .002) while average reach duration decreased (*β* = -0.177, *p* = .003). Hand-to-mouth frequency did not change significantly (*p* = .379), but average hand-to-mouth duration decreased (*β* = -1.583, *p* < .001). The frequency of vocalizations increased (*β* = 0.124, *p* = .002) while average vocalization duration decreased (*β* = -0.146, *p* = .011).

### 2.2. Behaviour-Specific Changes in Cortical Activity Across Age

Right-handed reaches were associated with a significant region of interest (ROI) x age interaction *F*(5,9718.49) = 7.99, p < .001, indicating that age-related changes in cortical activity differed by region. As shown in *Figure 2A* and detailed in *SI Appendix Table 2*, ROI-specific analyses revealed a significant decrease in bilateral posterior parietal cortex (PPC) activity (right: *β* = -0.026, *p* < .001; left: *β* = -0.026, *p* < .001). This indicates that as infants aged, their right-handed reaches were associated with decreased bilateral PPC activity.

**Figure 2.**
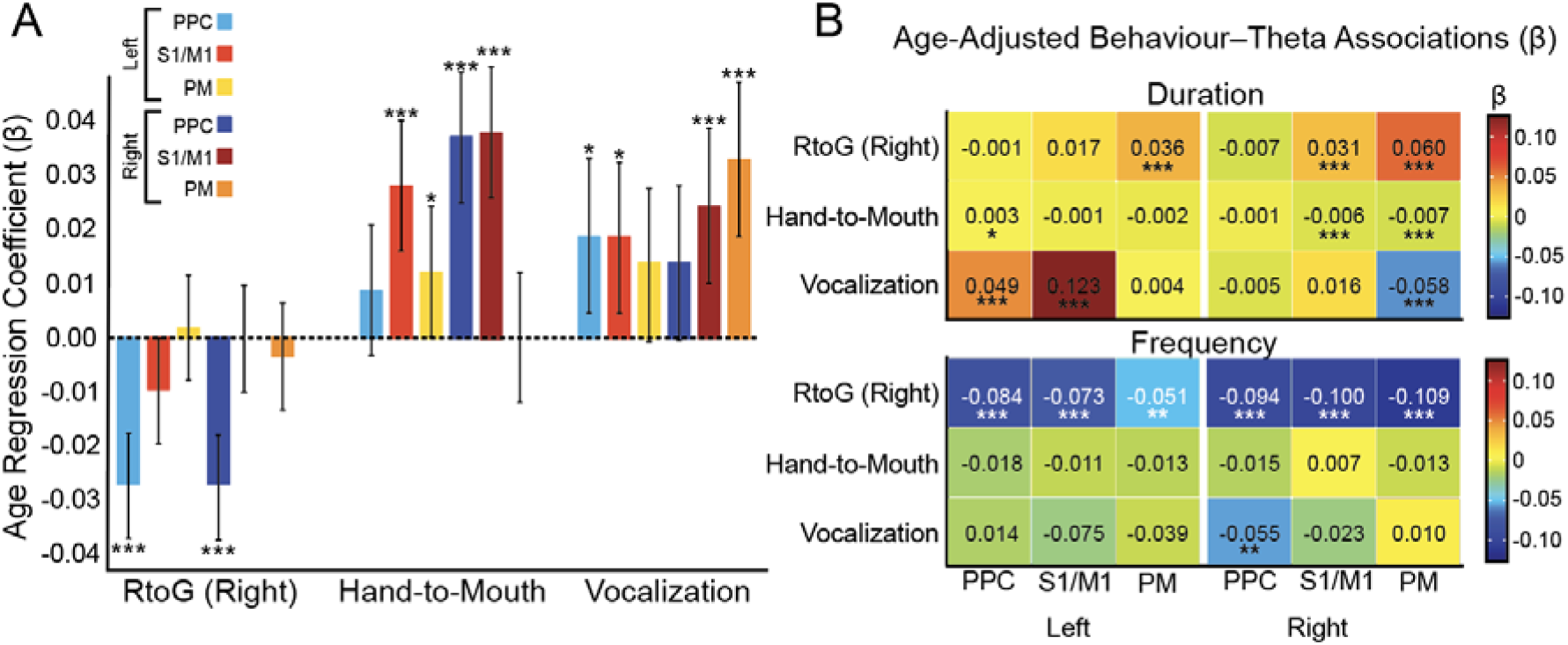
Panel A. Age-related changes in cortical activity during right-handed reach-to-grasp (left), hand-to-mouth (middle), and vocalization (right) behaviours. Age-related changes in cortical activity are shown for regions of interest (ROIs), including PPC, S1/M1, and PM. Error bars represent 95% CI and stars denote FDR corrected significance. *p* <.001***, *p* < .01**, *p* < .05*. Panel B. Heatmaps showing associations between behavioural variation and cortical activity after accounting for age. The top heatmap shows associations between the duration of reach-to-grasp, hand-to-mouth, and vocalization behaviours and corresponding cortical activity, whereas the bottom heatmap shows associations between behavioural frequency and cortical activity. Columns represent cortical regions of interest. Beta values indicate the direction of each association, with red representing positive and blue representing negative associations. Values within each cell are FDR corrected p-values. p <.001***, p < .01**, p < .05*. PPC = posterior parietal cortex; S1/M1 = somatomotor cortex; PM = premotor cortex.

Hand-to-mouth movements were also associated with a significant ROI x age interaction *F*(5,9359.19) = 8.14, *p* < .001, again indicating that age-related changes in cortical activation differed by region. Interestingly, with increasing age, hand-to-mouth movements were associated with increased activity in the bilateral somatomotor (S1/M1) region (right: *β* = 0.037, *p* < .001; left: *β* = 0.028, *p* < .001) as well as increased activation in the right PPC (*β* = 0.037, *p* < .001) and left premotor (PM) region (*β* = 0.013, *p* = .043).

Vocalizations in contrast, were not associated with a significant ROI x age interaction *F*(5,8309.41) = 1.20, *p* = .308, see *Figure 2A*, but were instead associated with a generalized increase in activity in all recorded regions as infants aged. After FDR correction, bilateral S1/M1 (right: *β* = 0.024, *p* < .001; left: *β* = 0.018, *p* = .016), right PM (*β* = 0.032, *p* < .001), and left PPC (*β* = 0.018, *p* = .016) activity all increased significantly during vocalizations as infants aged.

### 2.3. Behaviour-Specific Cortical Activity Independent of Age

After accounting for infant age, the relationship between movement duration and cortical activity differed across behaviours and cortical regions (*Figure 2B*). Interestingly, regardless of infant age, longer right-handed reach-to-grasp durations were associated with significantly increased right PM and S1/M1 activity, indicating that when infant reaches take longer and are less efficient, more ipsilateral cortical recruitment occurs. The pattern was reversed for hand-to-mouth movements; regardless of infant age, longer, more sustained hand-to-mouth bouts were associated with significantly less activity in right PM and S1/M1.

The relationship between behaviour frequency and cortical activity also differed by behaviour and cortical region after accounting for infant age (*Figure 2B*). As the frequency of right-handed reaches increased, cortical activity in all regions decreased whereas increased vocalization frequency was associated with less activity in the right PPC. For the full statistical analysis see *SI Appendix Table 3*.

## Discussion

This study introduces a novel data-driven approach for linking ongoing spontaneous behaviour to cortical activity in freely behaving infants without imposing repeated trials, strict baseline periods, or a prespecified HR shape. To our knowledge, this is the first application of FIR modelling to continuously coded spontaneous infant behaviour under naturalistic conditions. Using this approach, we identified age-related shortening of reach-to-grasp, hand-to-mouth, and vocalization behaviours; region-specific developmental changes in their associated cortical responses, and; stable relationships between cortical activity and both behavioural duration and behavioural frequency independent of infant age. These findings demonstrate that behaviour-specific cortical activation patterns can be resolved from the overlapping HRs of freely behaving infants, providing a novel and more ecologically valid, approach for testing theories of infant sensorimotor development under naturalistic conditions.

A common developmental pattern emerged across all three infant behaviours: reach-to-grasp, hand-to-mouth, and vocalization movements all became shorter with age, suggesting that they all become more discrete and efficiently organized. This interpretation is consistent with evidence that, as infants age, their reaches become faster and smoother, require fewer corrective movements, and increasingly anticipate object size and orientation (14, 15). Reach-to-grasp frequency, particularly for right-handed reaches, also increased with age, consistent with the developmental shift from a bimanual to right-handed reaching preference between 6 and 12 months (16). Hand-to-mouth movements showed a similar reduction in duration without a corresponding change in frequency, paralleling previous reports that this developmental period coincides with the introduction of solids and more functional feeding movements alongside a decline in non-nutritive sucking and exploratory hand-to-mouth movements (17). Finally, vocalizations, like reaches, became shorter but more frequent, consistent with previous reports of increasing vocal frequency, volubility, and complexity across infancy (18, 19). Together, these convergent patterns suggest that diverse infant behaviours become progressively more discrete, efficient, and controlled across development. Such coordinated nonlinear change is consistent with Dynamic Systems Theory, in which new behavioural organization emerges through interactions among developing neural, bodily, and environmental systems rather than through uniform maturation.

Infants generated increasingly efficient right-handed reach-to-grasp movements as they aged, which were accompanied by a bilateral reduction in PPC activity. Although the PPC is central to reaching, it also integrates sensory information for action planning and coordination of the eyes, limbs, and hands (20, 21). As infant reaching becomes more efficient, it relies less on trunk and whole-body postural support (22-24), potentially reducing the integrative demands on the PPC. Consistent with this interpretation, PPC activity decreases with short-term motor learning in adults (25), while infant motor development is associated with increasingly focal M1 activity and stronger frontoparietal coherence (26). Thus, as infants’ reaching becomes more efficient, the brain appears to accomplish the same goal with less reliance on PPC, suggesting increasing specialization and efficiency of the developing motor system. To our knowledge, the present data provide the first evidence that developmental increases in reaching efficiency are accompanied by reduced PPC recruitment in freely behaving infants.

Hand-to-mouth movements showed a contrasting developmental pattern. As infants aged, more discrete and efficient hand-to-mouth movements were associated with increasing bilateral S1/M1, right PPC, and left PM activity. Increased S1/M1 activity may reflect increasing attempts to coordinate hand-to-mouth movements by using tactile and proprioceptive feedback from the mouth to guide hand and finger movements (27-29). Greater PPC recruitment is also consistent with increasing demands to integrate oral somatosensory information with orofacial motor control during oral exploration, chewing, and feeding (30, 31), potentially accompanying the transition from non-nutritive sucking to more coordinated exploratory and nutritional hand-to-mouth movements. The age-related increase in left PM activity may further reflect greater demands on movement sequencing and hand-to-mouth coordination (32), consistent with the preferential recruitment of left PM during the learning and production of complex hand motor sequences in adults (32-35). Together, these findings suggest that hand-to-mouth development involves progressive reorganisation of sensorimotor networks to support the integration of somatosensory feedback with increasingly coordinated hand-to-mouth actions.

Vocalizations engaged a broader set of cortical regions than reach-to-grasp or hand-to-mouth movements, consistent with the greater sensorimotor complexity of emerging vocal control. In adults, increasingly demanding vocal sensorimotor transformations recruit bilateral somatomotor cortex (36) and children show greater somatomotor engagement than adults during speech motor learning (37). Thus, the age-related increase in somatomotor activity observed here may reflect increasingly complex and coordinated orofacial movements during vocal development. Similarly, increased ventral premotor cortex activity is thought to contribute to improved control of laryngeal, lingual, jaw, and lip movements required for vocal production (38, 39). The left PPC has been implicated in vocal sequencing and sensorimotor adaptation for speech (40, 41). Thus, increased left PPC activity with age may reflect growing demands to integrate sensory feedback with evolving vocal motor commands to correct and improve speech-like mouth movements. Together, these findings suggest that the early maturation of infant vocalizations is associated with increasing recruitment of distributed sensorimotor networks to support increasingly complex vocal control.

After accounting for age, variations in the frequency and duration of individual behaviours were associated with distinct patterns of cortical activity. Regardless of infant age, longer right-handed reach-to-grasp movements were associated with greater right S1/M1 and bilateral PM activity, consistent with evidence that increasing unimanual motor demands modulate M1 activity in adults (42) and engage closely interconnected somatomotor and premotor networks (43). In contrast, increasing reach-to-grasp frequency was associated with broadly reduced cortical activity, a pattern consistent with the TNGS prediction that repeated behaviour (reaching) promotes progressive refinement of motor and neuronal organization (1). Longer hand-to-mouth movements were associated with reduced right S1/M1 and PM activity, potentially reflecting sensorimotor adaptation or response attenuation during sustained non-nutritive sucking or oral exploration (31). Vocalization duration showed a different pattern, as longer vocalizations were associated with greater left PPC and S1/M1 activity but reduced right PM activity.

Electrocorticographic recordings have similarly revealed bilateral S1/M1 activity during infant babbling and cooing (44), whereas left PPC may help to integrate somatosensory feedback that drives corrective vocal adaptations during sustained phonation (45). Reduced right PM and PPC activity with increasing vocalization duration and frequency, respectively, may reflect decreasing motor demands when vocal behaviours are more sustained, although the functional significance of these effects remains uncertain. Together, these findings show that cortical recruitment varies with the specific frequency and duration of infants’ ongoing behaviours, revealing stable behaviour-specific cortical activity patterns that are not explained by age alone.

In sum, this study establishes a novel data-driven framework for linking spontaneous behaviours to corresponding cortical activity in human infants. By combining continuous behavioural coding with FIR modelling, we were able to exploit, rather than suppress, natural behavioural variability to resolve behaviour-specific HRs during reaching, hand-to-mouth, and vocalizations in human infants. Although these behaviours showed similar age-related changes in temporal structure (frequency and duration), each behaviour was associated with a distinct pattern of cortical development, and cortical activity remained related to behavioural variation even after accounting for age. To our knowledge, this is the first study to characterize cortical responses associated with multiple continuously occurring behaviours in freely behaving infants, providing a new framework for testing developmental theories, such as Dynamic Systems Theory, in more naturalistic and ecologically valid contexts. Future refinements that incorporate short-separation channels, broader cortical coverage, wider developmental sampling, and automated behavioural tracking will further improve the precision and scalability of this approach. Most importantly, these findings show that developmental changes in infant behaviour are accompanied by distinct, behaviour-specific reorganisations of cortical activity.

## Materials and Methods

### 4.1. Participants

A total of 59 infants (39 male, 20 female) aged 6-13 months (M = 9 months, S.D = 2.08) (205 sessions) were recruited from Kamloops, British Columbia. The study used a longitudinal design; participants were encouraged to come back monthly until they reached 14 months of age, thus each infant could potentially participate 7 times. We accepted infants for the first visit at any age between 6-13 months, leading to a total of 205 testing sessions (see *SI Appendix Table 4)*. After exclusions due to poor fNIRS signal, cap intolerance, or failure to finish the task, we had a total of 48 infants with 116 sessions. We grouped the infants and the corresponding sessions into four age groups: 6-7 months (27, M = 10; F = 17), 8-9 months (41, M = 13; F = 28), 10-11 months (28, M = 12; F = 16) and 12-13 months (20, M = 5; F = 15). All caregivers provided informed consent and information about birth date (expected and actual) and any known developmental conditions. Infants were eligible if they were born past 36 weeks, had no reported neurological or developmental disorders, and had no known hearing or visual impairments. The experiment was conducted at the Child Research Laboratory at Thompson Rivers University. The study and its procedures were approved by the Thompson Rivers University Human Research Ethics Board.

### 4.2. Experimental Task

The task ran for a maximum of 12 minutes, wherein the infants were encouraged to perform reach-to-grasp behaviour. In line with the naturalistic design of the experiment, the target was 12 reaches per session but was flexible based on the infant and caregiver. We aimed to collect 3 reaches in succession and accepted reaches to a variety of food items (cheerios, yogurt dots, smoothie puffs, or parent provided snacks) and if reaches to food items could not be obtained then we would switch to toys (lab-provided or caregiver-provided). Objects were placed on an open palm of either the caregiver or experimenter at the centre of the infant’s body roughly 2 inches away from their torso. After reaching, infants were encouraged to play and interact with experimenters and caregivers before the next reaching bout was initiated. Standardized videos were offered intermittently, but the infants continued to freely behave during these periods, and this was not differentiated in the continuous behavioural coding.

### 4.3. Data Collection

#### 4.3.1. Behavioural Data

Infants were fitted with neoprene fNIRS caps (46, 48 or 50; Artinis Medical Systems, Elst, The Netherlands) and fNIRS data acquisition was performed using OxySoft software (Artinis Medical Systems, Elst, The Netherlands). Two cameras captured frontal and midsagittal views of the participant, and E-prime software (E-Prime 3.0, Psychology Software Tools, 2020) was used to synchronize video and fNIRS recordings. Behavioural measures (*SI Appendix Table 5*) were manually coded using open-source software, BORIS (46).

#### 4.3.2. fNIRS Data Acquisition

Brain activity was measured using a continuous-wave fNIRS Brite system from Artinis (Artinis Medical Systems, Elst, The Netherlands). Appropriate infant head size (47) source-detector distances *(SI Appendix Table 6)* ranging from 20.6 mm to 41.2 mm (M = 28.4 mm) across 22 channels were used. Digitization was performed using a Polhemus 3D System (Patriot, Polhemus) registered to the Brodmann atlas to estimate the cortical sensitivity profile of each channel (*SI Appendix Figure 1C)* (48). Differential pathlength factor (DPF) values were calculated based on the infant’s age in years (0.5–1.08) and wavelength based on Scholkmann & Wolf, 2013 (49).

### 4.4. fNIRS Data Processing

The fNIRS data were processed using a custom application called FLARES (https://research.dezeeuw.ca/flares/), based on MNE-Python packages (https://doi.org/10.5281/zenodo.592483) (50). For full details of the methods see *Supplementary Methods and SI Appendix Figure 2*. Each channel was evaluated using several signal quality metrics and participants were removed entirely if 12 or more channels were identified as bad. Raw light intensity was converted to optical density and then processed through two signal filtering measures: temporal derivative distribution repair (TDDR) and wavelet filtering, before being converted to changes in oxyhemoglobin (HbO) and deoxyhemoglobin (HbR) concentrations using the modified Beer–Lambert law (51, 52). Finally, the hemoglobin concentrations were passed through both frequency and bandwidth filters before events were extracted from the coded behavioural file from BORIS and added to the design matrix for the FIR model. For all parameters selected for fNIRS data processing in the FLARES application, see *SI Appendix Table 7*.

### 4.6. Statistical Analyses

Three linear mixed-effects models were fitted to examine: 1) behavioural development, 2) age-related changes in behaviour-specific cortical activation, and 3) stable brain-behaviour associations for the three target behaviours (right-handed reaches, hand-to-mouth, and vocalizations) independent of age. Participant was included as a random intercept in all models. Model-specific fixed effects included age, ROI, behavioural measures (frequency or duration), and interactions where appropriate. In the third brain-behaviour model age was included as a covariate. Because the timing of the HR peak varied across behaviours, participants, or ROIs, no single FIR time bin consistently captured the peak response. Therefore, FIR bins were included as regressors in the fNIRS analyses. This allows the model to estimate differences across the HR window rather than selecting an arbitrary peak time point. False discovery rate (FDR) correction was applied across all comparisons for each model. Model formulas and further details can be found in *Supplementary Methods*.

## Acknowledgments

We would like to thank and acknowledge the research participants and their caregivers for their contributions to the study. We also thank Supreeta Ranchod, Avory Zienowicz, Liesl Milovick, and Disha Datta for their assistance with data collection. This research was funded by the support of the New Frontiers in Research Fund (JMK), [NFRFRE-2022-00173], the Natural Sciences and Engineering Research Council of Canada (JMK), [RGPIN-2017-05995], and the Thompson Rivers University Undergraduate Research Experience Award Program (UREAP).

## Notes

### Competing Interest Statement

The authors have declared no competing interest.

