## Supporting Information Appendix for "Distinct Patterns of Cortical Reorganisation Accompany Behaviour-Specific Development in Freely Behaving Infants"

**Results**

**Table 1.** Results from linear mixed-effects models on reach-to-grasp, hand-to-mouth, and vocalization behaviour frequency and average duration as an effect of age. Reported significance values are FDR-corrected.

| **Behaviour** | **Frequency (events/min)** | **Average Duration (s)** |
| --- | --- | --- |
| **Reach-to-grasp (Right)** | *β* = 0.087, SE = 0.028, *t*(109.98) = 3.45, *p* = .002. ** | *β* = -0.177, SE = 0.056, *t*(101.46) = -3.16, *p* = .003. ** |
| **Hand-to-Mouth** | *β* = 0.044, SE = 0.049, *t*(116.24) = 0.88, *p* = .379. | *β* = -1.583, SE = 0.372, *t*(87.43) = -4.251, *p* < .001. *** |
| **Vocalization** | *β* = 0.124, SE = 0.037, *t*(116.92) = 3.314, *p* = .002. ** | *β* = -0.146, SE = 0.055, *t*(93.82) = -2.649, *p* = .011. * |

**Table 2**. Results from linear mixed-effects model for ROI x age follow-up interactions. p <.001***, p < .01**, p < .05*.

| **Behaviour** | **Hemisphere** | **ROI** | **β** | **95% CI** | **FDR-corrected p value** |
| --- | --- | --- | --- | --- | --- |
| Reach-to Grasp  (Right Hand) | Right | Posterior Parietal | -0.026 | -0.036 – -0.016 | < .001*** |
|  |  | Somatomotor | 0.001 | -0.009 – 0.011 | 0.820 |
|  |  | Premotor | -0.003 | -0.013 – 0.007 | 0.671 |
|  | Left | Posterior Parietal | -0.026 | -0.036 – -0.016 | < .001*** |
|  |  | Somatomotor | -0.009 | -0.019 – 0.000 | 0.095 |
|  |  | Premotor | 0.002 | -0.007 – 0.012 | 0.696 |
| Hand-to-Mouth | Right | Posterior Parietal | 0.037 | 0.026 – 0.048 | < .001*** |
|  |  | Somatomotor | 0.037 | 0.026 – 0.048 | < .001*** |
|  |  | Premotor | 0.000 | -0.010 – 0.013 | 0.820 |
|  | Left | Posterior Parietal | 0.010 | -0.002 – 0.021 | 0.111 |
|  |  | Somatomotor | 0.028 | 0.017 – 0.039 | < .001*** |
|  |  | Premotor | 0.013 | 0.002 – 0.024 | 0.043* |
| Vocalization | Right | Posterior Parietal | 0.014 | 0.001 – 0.028 | 0.057 |
|  |  | Somatomotor | 0.024 | 0.011 – 0.037 | < .001*** |
|  |  | Premotor | 0.032 | 0.019 – 0.046 | < .001*** |
|  | Left | Posterior Parietal | 0.018 | 0.005 – 0.032 | 0.016* |
|  |  | Somatomotor | 0.018 | 0.005 – 0.031 | 0.016* |
|  |  | Premotor | 0.014 | 0.000 – 0.027 | 0.063 |

**Table 3.** Associations between cortical activity and spontaneous behaviour. Results from linear mixed-effects models for duration and frequency, with theta as the dependent variable, age (centred) as a covariate, region of interest (ROI), the behavioural predictor, and the ROI x behaviour interaction as fixed effects. Significance values reported are FDR-corrected. *p* <.001***, *p* < .01**, *p* < .05*.

| **Duration** | | | | | |
| --- | --- | --- | --- | --- | --- |
| **Behaviour** | **Hemisphere** | **ROI** | **β** | **95% CI** | **FDR-corrected p value** |
| Reach-to Grasp  (Right Hand) | Right | Posterior Parietal | -0.007 | -0.023 – 0.009 | 0.493 |
|  |  | Somatomotor | 0.031 | 0.015 – 0.047 | < .001*** |
|  |  | Premotor | 0.060 | 0.044 – 0.076 | < .001*** |
|  | Left | Posterior Parietal | 0.001 | -0.015 – 0.017 | 0.867 |
|  |  | Somatomotor | 0.017 | 0.000 – 0.032 | 0.091 |
|  |  | Premotor | 0.036 | 0.020 – 0.052 | < .001*** |
| Hand-to-Mouth | Right | Posterior Parietal | -0.001 | -0.003 – 0.002 | 0.729 |
|  |  | Somatomotor | -0.006 | -0.008 – -0.003 | < .001*** |
|  |  | Premotor | -0.007 | -0.010 – -0.004 | < .001*** |
|  | Left | Posterior Parietal | 0.003 | 0.000 – 0.006 | 0.031* |
|  |  | Somatomotor | -0.001 | -0.004 – 0.002 | 0.515 |
|  |  | Premotor | -0.002 | -0.005 – 0.000 | 0.261 |
| Vocalization | Right | Posterior Parietal | -0.005 | -0.031 – 0.020 | 0.729 |
|  |  | Somatomotor | 0.016 | -0.009 – 0.042 | 0.323 |
|  |  | Premotor | -0.058 | -0.083 – -0.032 | < .001*** |
|  | Left | Posterior Parietal | 0.049 | 0.023 – 0.075 | < .001*** |
|  |  | Somatomotor | 0.123 | 0.097 – 0.148 | < .001*** |
|  |  | Premotor | 0.004 | -0.022 – 0.030 | 0.785 |
| **Rate** | | | | | |
| Reach-to Grasp  (Right Hand) | Right | Posterior Parietal | -0.094 | -0.127 – -0.061 | < .001*** |
|  |  | Somatomotor | -0.100 | -0.133 – -0.067 | < .001*** |
|  |  | Premotor | -0.109 | -0.143 – -0.076 | < .001*** |
|  | Left | Posterior Parietal | -0.084 | -0.117 – -0.051 | < .001*** |
|  |  | Somatomotor | -0.073 | -0.107 – -0.039 | < .001*** |
|  |  | Premotor | -0.051 | -0.084 – -0.018 | 0.006** |
| Hand-to-Mouth | Right | Posterior Parietal | -0.015 | -0.036 – 0.006 | 0.282 |
|  |  | Somatomotor | 0.007 | -0.014 – 0.027 | 0.637 |
|  |  | Premotor | -0.009 | -0.030 – 0.011 | 0.493 |
|  | Left | Posterior Parietal | -0.018 | -0.039 – 0.002 | 0.162 |
|  |  | Somatomotor | -0.011 | -0.032 – 0.009 | 0.407 |
|  |  | Premotor | -0.013 | -0.034 – 0.008 | 0.323 |
| Vocalization | Right | Posterior Parietal | -0.055 | -0.090 – -0.021 | 0.004** |
|  |  | Somatomotor | -0.023 | -0.057 – 0.012 | 0.323 |
|  |  | Premotor | 0.010 | -0.024 – 0.044 | 0.648 |
|  | Left | Posterior Parietal | -0.016 | -0.050 – 0.017 | 0.473 |
|  |  | Somatomotor | -0.023 | -0.057 – 0.012 | 0.232 |
|  |  | Premotor | -0.010 | -0.044 – 0.024 | 0.648 |

**Materials and Methods**

**4.1. Participants**

The participants were recruited from surrounding areas through a combination of online advertising, posters, and engagement with community connections (childcare services, social services, parent groups, and healthcare facilities). Caregivers received a $40 gift card as compensation for each testing session. Infants could only participate in one session per month, and repeated observations were handled with participant as a random intercept. As shown in Table 4, after exclusions the final number of infants was 48 with 116 sessions remaining.

**Table 4.** Participant information and exclusion. ‘Did not complete task’ denotes any child who would not participate for a minimum of 10 minutes. Cap intolerance denotes any infant who would not wear the cap or removed the cap mid-session. Poor fNIRS signal denotes any session that was excluded for having more than 12 channels that failed quality control metrics in fNIRS processing. Other/missing data involved missing files. The infants (n) column indicates the total number of infants that had one session of the denoted files missing. An infant missing one of their sessions was not excluded from the analysis completely unless all sessions were removed/missing for any reason.

|  | **Infants (n)** | **Session (n)** | **Sessions excluded (%)** |
| --- | --- | --- | --- |
| **Total participated** | 59 | 205 |  |
| **Did not complete task** | 3 | 3 | 1% |
| **Cap intolerance** | 12 | 21 | 10% |
| **Poor fNIRS signal** | 35 | 63 | 31% |
| **Other/missing data** | 2 | 2 | 1% |

**4.2. Data Collection**

*4.2.1. Behavioural Data Acquisition and Scoring*

Infants were habituated to the testing environment while experimenters took head measurements to identify and mark Central zero (Cz). The infants were then placed in an infant chair on the floor (*Figure 3A*) and fitted with one of three neoprene fNIRS caps (46, 48 or 50; Artinis Medical Systems, Elst, The Netherlands). fNIRS data acquisition was performed using OxySoft software (Artinis Medical Systems, Elst, The Netherlands). Once the cap was fitted, the experimenters started two cameras (Panasonic Lumix GH6, 59.94 frames per second) which captured frontal and midsagittal views of the participant *(SI Appendix Figure 1A).* The trial started when the fNIRS acquisition computer were displayed for the cameras and a flash on the screen was used to sync the video and fNIRS data, using E-Prime (E-Prime 3.0, Psychology Software Tools, 2020).

**Table 5.** The behavioural scoring parameters for infant movements. Starred behaviours denote target behaviours for analysis and the remaining behaviours were used as nuisance regressors in the FIR model.

| **Behaviour** | **Description** |
| --- | --- |
| Reach-to-Grasp | Any time the infant reaches their arm (either right, left, or both) to grasp an object (toy, food, etc.). The reach-to-grasp movement starts at the lift stage (time point at which the hand made its first discernable movement toward the object) and is completed when the infant was holding the object (time point at which digit closure was complete, resulting in successful holding of the target) as defined by (1). |
| Kick | Any leg movements (either right, left, or both) that differed significantly from the baseline seated position. Movement starts when the foot lifts off the floor and stops when the infant returns to their baseline (relaxed) leg state. |
| Hand-to-Mouth | The infant places an item in their mouth; this could be their own hand, food, individual digits, or a toy. The movement started from the frame in which the item touched the mouth and ended when either the food object was consumed (chewing stopped) or when they withdrew the object or hand. |
| Vocalization | Any sound the infant emitted was scored as a vocalization. The event started when the utterance started and was scored as completed when the utterance ceased. In cases of babbling with several short bursts of vocalizations in a string, this was coded as one vocal event. For example, a yell (yaaaaaaaaa) and a babble (ba, ba, ba, ba, ba) were both scored as one event. |
| Extraneous Movement | Any movement that lacked intention or a clear directive was coded as extraneous movement, so the FIR model could still model movement. Many of these movements would be spastic benign myoclonus-like movements (2). The movement started when any significant deviation from the infants’ baseline state was made that was not one of the other scored behaviours. |
| Self-Touch | Any instance where punctuate contacts by the hand with the body were subject to analysis. All types of hand contacts (rostral or caudal body contacts) and hand posture (dorsum, palmar, or grasp contacts) (3) were combined into one measure of self-touch. This was not sub-divided by hand. |
| Dancing | Repetitive movements (≥2 repetitions) that were rhythmic in nature. This could include head-bobbing synchronously with leg or arm movements, whole-body swaying, kicking with body movements, or any other combination. |
| Head Movement | Any instance where the infant independently moves their head more than 45 degrees from the baseline (forward facing) posture. This movement could be side-to-side or up-and-down, and encompasses turns, tilts, nods, or shakes that could not be attributed to whole-body displacement. |
| Throw | Instances where the infant has an object in hand (any object including food items) and raises their arm to release the object with intentional velocity. If the infant simply raised the arm and the object was dropped this was not coded as a throw. |

*4.2.2. fNIRS Data Acquisition*

Brain activity was measured using a continuous-wave fNIRS Brite system from Artinis (Artinis Medical Systems, Elst, The Netherlands). The system uses two wavelengths of near-infrared light (762 nm and 842 nm) and source power is automatically controlled through multi-power gain settings which operate within international safety standards (IEC-60825-1). Data were sampled at 25 Hz using 10 source and 8 detector optodes arranged bilaterally to form 22 measurement channels (11 per hemisphere). The optodes were placed within one of three (46, 48, and 50 cm) wearable neoprene caps to optimize scalp coupling and signal quality (Artinis Medical Systems, Elst, The Netherlands).

Probe geometry (*SI Appendix Figure 1B*) was the same across cap sizes, to preserve relative spacing of sources and detectors. Optode placement was standardized using cranial landmarks (nasion, inion, and preauricular points).

**Table 6.** The distances in millimetres between each source and detector for each cap. Channels shorter than 15 mm and longer than 45 mm were automatically removed during preprocessing.

| Source Number | Detector Number | Channel ID | 46cm cap | 48cm cap | 50cm cap |
| --- | --- | --- | --- | --- | --- |
| 1 | 1 | 1 | 28.0 | 24.6 | 33.3 |
| 2 | 1 | 2 | 29.3 | 23.5 | 31.8 |
| 1 | 2 | 3 | 36.0 | 34.5 | 37.7 |
| 2 | 2 | 4 | 28.8 | 29.1 | 42.8 |
| 3 | 2 | 5 | 27.6 | 23.3 | 25.8 |
| 2 | 3 | 6 | 38.0 | 27.7 | 32.4 |
| 4 | 3 | 7 | 25.5 | 28.3 | 30.1 |
| 2 | 4 | 8 | 29.0 | 23.2 | 27.1 |
| 3 | 4 | 9 | 28.1 | 22.4 | 37.2 |
| 4 | 4 | 10 | 25.6 | 31.1 | 32.3 |
| 5 | 4 | 11 | 25.1 | 35.0 | 29.5 |
| 6 | 5 | 12 | 28.7 | 23.7 | 31.4 |
| 7 | 5 | 13 | 28.3 | 23.2 | 25.8 |
| 6 | 6 | 14 | 36.5 | 31.3 | 30.1 |
| 7 | 6 | 15 | 28.5 | 32.8 | 33.9 |
| 8 | 6 | 16 | 35.2 | 27.9 | 30.5 |
| 7 | 7 | 17 | 38.2 | 31.7 | 30.4 |
| 9 | 7 | 18 | 19.9 | 32.3 | 23.4 |
| 7 | 8 | 19 | 30.0 | 29.6 | 23.9 |
| 8 | 8 | 20 | 30.7 | 32.7 | 23.6 |
| 9 | 8 | 21 | 22.9 | 30.7 | 25.7 |
| 10 | 8 | 22 | 29.1 | 32.4 | 29.0 |

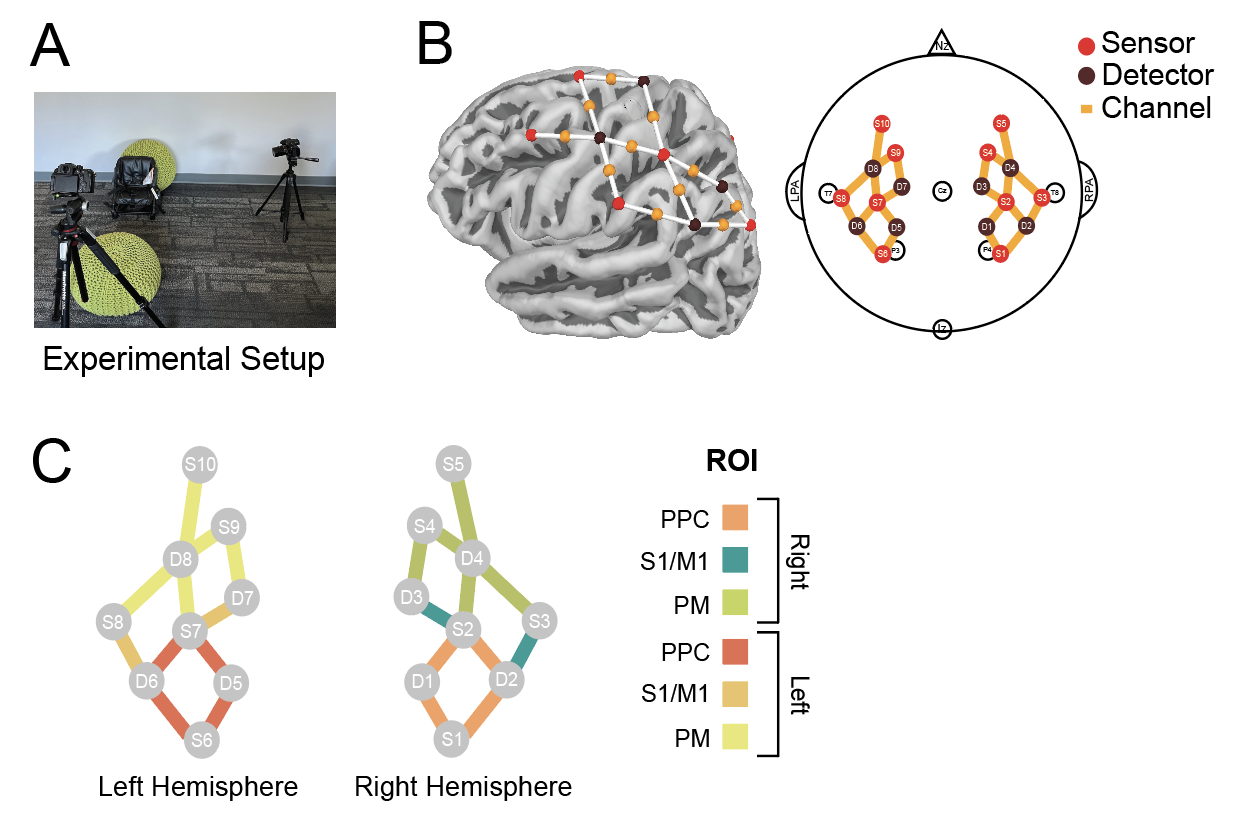

**Figure 1.** Panel A shows the experimental set up along with the camera positions. Panel B shows the sources (red), detectors (brown), and corresponding channels (yellow) on the left hemisphere of a 3D brain created using Polhemus digitization. Beside the brain rendering there is a bilateral display of the optode placement overlaid on a 2D head with key 10-20 points highlighted. Panel C shows the optode array with colour-coded lines indicating the region of interest and hemisphere the channels belong to.

**4.3. fNIRS Data Processing**

FLARES was first used to format the raw fNIRS data file (SNIRF file) by adding behavioural markers, time-synchronizing the fNIRS and behavioural data, and updating the optode array into a 3D array. To add the behavioural markers, we added the behaviourally coded events from BORIS and integrated those coded events into the SNIRF file, using our time sync data to ensure these events were appropriately time-locked. Next, we updated the optode array imported from OxySoft, which exists in a 2D format, into a 3D array using the files created with Polhemus digitization based on the cap size the infant was wearing for each session. At this point, the updated SNIRF file was loaded into the FLARES application. Once the optode positions had been verified (that they were successfully transferred into 3D points), pre-processing could start.

Each channel passed through several signal quality metrics: scalp coupling index, peak spectral power, signal-to-noise ratio, coefficient of variation, median absolute deviation and power spectral density noise (4-8). Raw light intensity was converted to optical density and then processed using two signal-filtering measures: Temporal Derivative Distribution Repair and wavelet filtering before being converted to changes in oxyhemoglobin (HbO) and deoxyhemoglobin (HbR) concentrations using the modified Beer-Lambert law (9, 10). Finally, the hemoglobin concentrations were passed through both frequency and bandwidth filters before events were extracted from the coded behavioural file and added to the design matrix for the FIR model.

The FIR model estimated the hemodynamic response using 15 consecutive 1-second bins spanning a total of 15 seconds following behavioural event onset. For duration modelling, every behaviour was treated as an onset event regardless of its duration. Concurrent behaviours were entered as separate regressors within the same design matrix, allowing the FIR coefficient for each target behaviour to be estimated while also accounting for overlapping behaviours. All behaviours in Table 5, excluding right-handed reach-to-grasp, hand-to-mouth, and vocalizations were used as nuisance regressors.

**Figure 2.** fNIRS processing methods performed in the FLARES application.

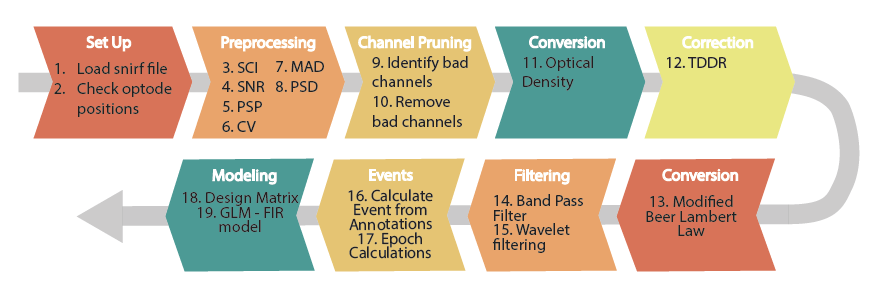

**Table 7**. FLARES output parameters.

| **Category** | **Parameter** | **Selection** |
| --- | --- | --- |
| Preprocessing | Down sample | True |
|  | Down sample frequency | 25 Hz |
| Trimming | Trim | True |
|  | Seconds to Keep | 5.0 |
| Short/Long Channels | Short Channels | False |
|  | Short Channel Threshold | 15 mm |
|  | Long Channel Threshold | 45 mm |
| Heart Rate | Heart Rate | True |
|  | Seconds to strip HR | 5.0 seconds |
|  | Low heartrate | 40 bpm |
|  | High heartrate | 200 bpm |
|  | Smoothing window | 100 seconds |
|  | Heart rate window | 25 seconds |
| Scalp Coupling Index (SCI) | SCI | True |
|  | SCI time window | 3.0 seconds |
|  | SCI threshold | 0.6 |
| Signal to Noise Ratio (SNR) | SNR | True |
|  | SNR threshold | 5.0 |
| Peak Spectral Power (PSP) | PSP | True |
|  | PSP time window | 3.0 seconds |
|  | PSP threshold | 0.1 |
| Coefficient of Variation (CV) | Coefficient of Variation | True |
|  | Coeff Threshold | 20 % noise threshold |
| Median Absolute Deviation (MAD) | MAD | True |
|  | MAD Threshold | 4 deviations |
| Power Spectral Density Noise (PSD) | PSD Noise | True |
|  | Target frequency division | 4 divisions |
|  | dB Limit | -60 dB |
| Sensor Dropout | Sensor Dropout | True |
|  | Sensor Dropout Variance Threshold | 0.05 % |
| Bad Channel Handling | Bad Channel Handling | Remove |
|  | Max Bad Channels | 24 (oxy and deoxy) 12 channels total |
| Temporal Derivative Distribution Repair Filtering (TDDR) | TDDR | True |
| Wavelet Filtering | Wavelet Filtering | True |
|  | IQR | 1.5 |
|  | Wavelet Type | Db4 |
|  | Wavelet Level | 3 |
| Filtering | Filtering | True |
|  | Low frequency | 0.005 |
|  | High frequency | 0.3 |
|  | Low transitional bandwidth | 0.002 |
|  | High transitional bandwidth | 0.002 |

**4.4. ROI construction**

To determine channel-level anatomical localization we first digitized each cap using a Polhemus 3D System (Patriot, Polhemus Inc., Colchester, VT, USA). Due to practical challenges associated with digitizing optode positions on infants, digitization was performed on three mannequin heads sized appropriately for each cap. One digitization template was obtained for each cap size and applied to all sessions using that cap size. To estimate the cortical regions associated with each measurement channel we used fNIRS Optodes’ Location Decider (fOLD) (11) packaged within FLARES. Optode positions were registered to the Brodmann atlas to estimate the cortical sensitivity profile of each channel.

Channels were then grouped into three ROIs (see supplemental Table 8) within each hemisphere, posterior parietal cortex (PPC), somatomotor cortex (S1/M1), and premotor cortex (PM) areas. Channels were assigned to an ROI when they had over 38% coverage within the Brodmann areas (BA) defining the ROI. The PPC was defined by BA 39 and BA 40 (12), S1/M1 by BA 2, BA 3, and BA 4, which overlaps primary somatosensory and primary motor cortices, and PM by BA 6 (13). Although individual channels were sensitive to adjacent Brodmann areas, these secondary contributions were not used to define the ROIs. The channel-level theta estimates obtained from the GLM were averaged across channels within each ROI, to produce ROI-level theta estimates. When a channel was removed at the signal quality metric assessment stage in FLARES, the remaining channels available were used to complete the average.

**Table 8.** Region of Interest (ROI) Breakdown.

| **Hemisphere** | **ROI** | **Defining Brodmann Areas** | **Channels Included** |
| --- | --- | --- | --- |
| Right | PPC | BA 39/40 | S1_D1; S1_D2; S2_D1; S2_D2 |
|  | S1/M1 | BA 2,3 & 4 | S2_D3; S3_D2 |
|  | PM | BA 6 | S2_D4; S3_D4; S4_D3; S4_D4; S5_D4 |
| Left | PPC | BA 39/40 | S6_D5; S6_D6; S7_D5; S7_D6 |
|  | S1/M1 | BA 2,3 & 4 | S7_D7; S8_D6 |
|  | PM | BA 6 | S7_D8; S8_D8; S9_D7; S9_D8; S10_D8 |

**4.5. Statistical Analyses**

Statistical analyses were conducted in R using the lme4 (14), lmerTest (15), and emmeans (16) packages. Linear mixed-effects models were fitted using lme4 and lmerTest, with Satterthwaite-approximated degrees of freedom to evaluate statistical significance of fixed effects. Sum-to-zero contrasts were applied to ROI, so no ROI was treated as a reference category, and ROI-specific slopes were estimated using emtrends from the emmeans package. Multiple comparisons were controlled using the Benjamini-Hochberg false discovery rate (FDR).

We analyzed the three target behaviours, right-handed reach-to-grasp actions, hand-to-mouth, and vocalizations, using average duration and behavioural frequency (events/min). A linear mixed-effects model was used for each behaviour and dependent variable (events/min and duration), with age in months as a continuous variable and participant as a random intercept. The significance of fixed effects was evaluated using Satterthwaite-approximated degrees of freedom, and FDR correction was applied across the six behavioural age analyses. HbO and HbR were both collected but only HbO was used to perform the final analysis because HbO has been shown to have higher reliability and stronger task sensitivity (17-19).

To understand the developmental trajectory of cortical activity associated with spontaneous behaviour, we examined the age-related trajectories of the fNIRS response (theta values) separately for each target behaviour (reach-to-grasp, hand-to-mouth, and vocalizations). A linear mixed-effects model was used including age, region of interest (ROI), and the ROI x age interaction. FIR bins were included as a categorical fixed-effect covariate to account for variation in theta across the time course of the hemodynamic response. FDR correction was applied across all ROI comparisons.

To understand the relationship between cortical activity and spontaneous behaviour we examined the fNIRS response separately for duration and rate of each target behaviour (reach-to-grasp, hand-to-mouth, and vocalizations). Separate linear mixed-effects models were used for behavioural duration and rate, with theta as the dependent variable. In each model, age (centred) was included as a covariate. Region of interest (ROI), the behavioural predictor, and the ROI x behaviour interaction were included as fixed effects. The FIR bins were included as nuisance regressors to account for the variability in the peak of the hemodynamic response and participant was included as a random intercept. FDR correction was applied across all 36 ROI x behaviour comparisons.
